# Single Agent and Synergistic Activity of Maritoclax with ABT-263 in Nasopharyngeal Carcinoma (NPC) Cell Lines

**DOI:** 10.1101/471888

**Authors:** Benedict Shi Xiang Lian, Kwok-Wai Lo, Alan Soo-Beng Khoo, Nethia Mohana-Kumaran

**Affiliations:** School of Biological Sciences, Universiti Sains Malaysia, Penang, Malaysia; Department of Anatomical & Cellular Pathology, The Chinese University of Hong Kong Hong Kong SAR; Molecular Pathology Unit, Cancer Research Centre, Institute for Medical Research, Jalan Pahang, Kuala Lumpur, Malaysia

**Keywords:** Nasopharyngeal carcinoma, 3D NPC spheroids, Maritoclax, ABT-263, BH3 mimetics

## Abstract

**Objective:** The BCL-2 anti-apoptotic proteins are over-expressed in many cancers and hence are attractive therapeutic targets. In this study we tested the sensitivity of two NPC cell lines HK1 and C6661-1 to Maritoclax which is reported to repress anti-apoptotic protein MCL-1 and BH3 mimetic ABT-263 which selectively inhibits anti-apoptotic proteins BCL-2, BCL-XL and BCL-w. We investigated the sensitization of the NPC cell lines to these drugs using the SYBR Green I assay and 3D NPC spheroids.

**Results:** We report that Maritoclax repressed MCL-1, BCL-2, and BCL-XL in a dose- and time-dependent manner and displayed a single agent activity in inhibiting cell proliferation of the NPC cell lines. Moreover, combination of Maritoclax and ABT-263 exhibited synergistic cell proliferation effect in the HK1 cell line. Similar results were obtained in the 3D spheroids. More notably, 3D spheroids either treated with single agent Maritoclax or combination with ABT-263 over 10 days did not develop resistance to the treatment rapidly. Collectively, the findings illustrate that Maritoclax as a single agent or combination with BH3 mimetics could be a potential treatment strategy for NPC but further studies in preclinical models are warranted to fully unravel the prospects of these drugs.

## Introduction

NPC is the sixth most common cancer in Southeast Asia (SEA) with Malaysia reporting one of the highest national incidences in SEA [1]. Treating patients with metastatic NPC is often a challenge as patients develop resistance to systemic anti-cancer therapies such as chemotherapy and retreating local recurrence with radiotherapy have many limitations. Thus, novel or improved treatment strategies are urgently needed to curb this disease.

The BCL-2 family proteins are critical regulators of the intrinsic apoptosis pathway [2]. The anti-apoptotic BCL-2 proteins are reported to be overexpressed in many cancers and thus are attractive therapeutic targets. Utilizing the immunohistochemistry (IHC) technique, BCL-2 expression was detected in 80% NPC tissues and 71% adjacent dysplastic lesions compared to normal nasopharynx epithelia [3]. Employing the same technique, similar results were obtained in other studies [4-6]. NPC tissues which were positive for BCL-2 expression, highly correlated with neck lymph nodes metastasis [4] and a worse disease-free 5-year survival [7]. Collectively, these studies suggest that the BCL-2 anti-apoptotic proteins are relevant targets for NPC treatment.

In the present study, we investigated the sensitivity of two NPC cell lines HK1 and C666-1 to single agent treatment of Maritoclax and ABT-263 alone and in combination using 2-dimensional (2D) and 3-dimensional (3D) cell culture models. ABT-263 binds with high affinity to BCL-2 and BCL-XL [8] whereas Maritoclax was reported to antagonize MCL-1 and target the protein for proteasome-mediated degradation [9].

## Methods

### Cells and Cell culture

The human NPC cell lines HK1 and C666-1 were grown and authenticated using the AmpFISTR profiling as described [10]. Cells were not passaged unnecessarily and experiments were performed within 2-3 passages of the foundation stocks.

### Western blot analysis

Cells were lysed and analyzed on reducing SDS-PAGE as described [11]. Membranes were blocked with 5% blotto and probed with antibodies against: MCL-1 (clone 22, BD Pharmingen, USA), BCL-2 (clone 100, BD Pharmingen, USA), BCL-XL (clone 2H12, BD Pharmingen, USA) and α-tubulin (Y1/2, Abcam, UK). Membranes were washed with PBST several times and bound antibodies were detected using either goat anti-mouse IgG-HRP (sc-2005, Santa Cruz Biotechnology, USA) or goat anti-rat IgG-HRP (sc-2006, Santa Cruz Biotechnology, USA) and enhanced with Super Signal^®^ West Pico Chemiluminescent (Thermo Scientific, USA).

### SYBR Green I assay and Synergy analysis

The SYBR Green I assay was conducted as described [12, 13]. The HK1 cells were seeded at a density of 2500 cell/well and the C666-1 cells were seeded at a density of 4000 cells/well in 96-well plates and left to attach for 6-7 hours. Cells were treated to a concentration series [0-32 μM of single agent Maritoclax (ChemScene, USA) and ABT-263 (Selleckchem, USA) and as combination along the long plate axis for 72 hours. Sensitization to ABT-263 by Maritoclax was assessed by testing a fixed concentration of Maritoclax to increasing concentrations and ABT-263 for 72 hours. Cell proliferation was quantified as described [14]. Synergy studies were conducted by calculating the combination index (CI) using the CalcuSyn 2.11 software (Biosoft Inc, Cambridge, UK). A CI of <1 indicates synergistic effect, CI =1 an additive effect, and CI > 1 antagonism.

### Generation of 3D spheroids

Approximately 5000 (2.5 × 10^4^ / ml) HK1 cells were seeded in the ultra-low attachment (ULA) 96-well U bottom plate (Corning, USA) and centrifuged at 1200 rpm for 2 minutes. The plates were incubated in a humidified incubator at 37°C with 5% CO_2_ for three days. Spheroids were embedded in collagen matrix as described [15, 16]. Spheroids were treated with ABT-263 and Maritoclax at doses and times required in 1 ml of complete medium and were incubated in a humidified incubator at 37°C with 5% CO_2_. Phase contrast snapshots of spheroids were taken every 24 hours using the IX71 Olympus inverted fluorescence microscope over 10 days to document spheroid growth and invasion. Spheroids were washed 3 times in PBS and stained with 4 mmol/L calcein AM and 2 mmol/L ethidium homodimer I (Thermo Fisher Scientific, USA) for an hour at 37°C. Images of spheroids were captured using the IX71 Olympus inverted microscope.

## Results

### Maritoclax suppresses BCL-2 anti-apoptotic proteins in a dose- and time-dependent manner

The basal expression levels of the anti-apoptotic proteins MCL-1, BCL-XL and BCL-2 in the NPC cells were first investigated. MCL-1 was expressed in both HK1 and C666-1 cells (Fig. 1a). Higher expression of BCL-2 was found in the C666-1 cells, whereas relatively higher expression of BCL-XL was found in the HK1 cells (Fig. 1a).

**Figure 1.**
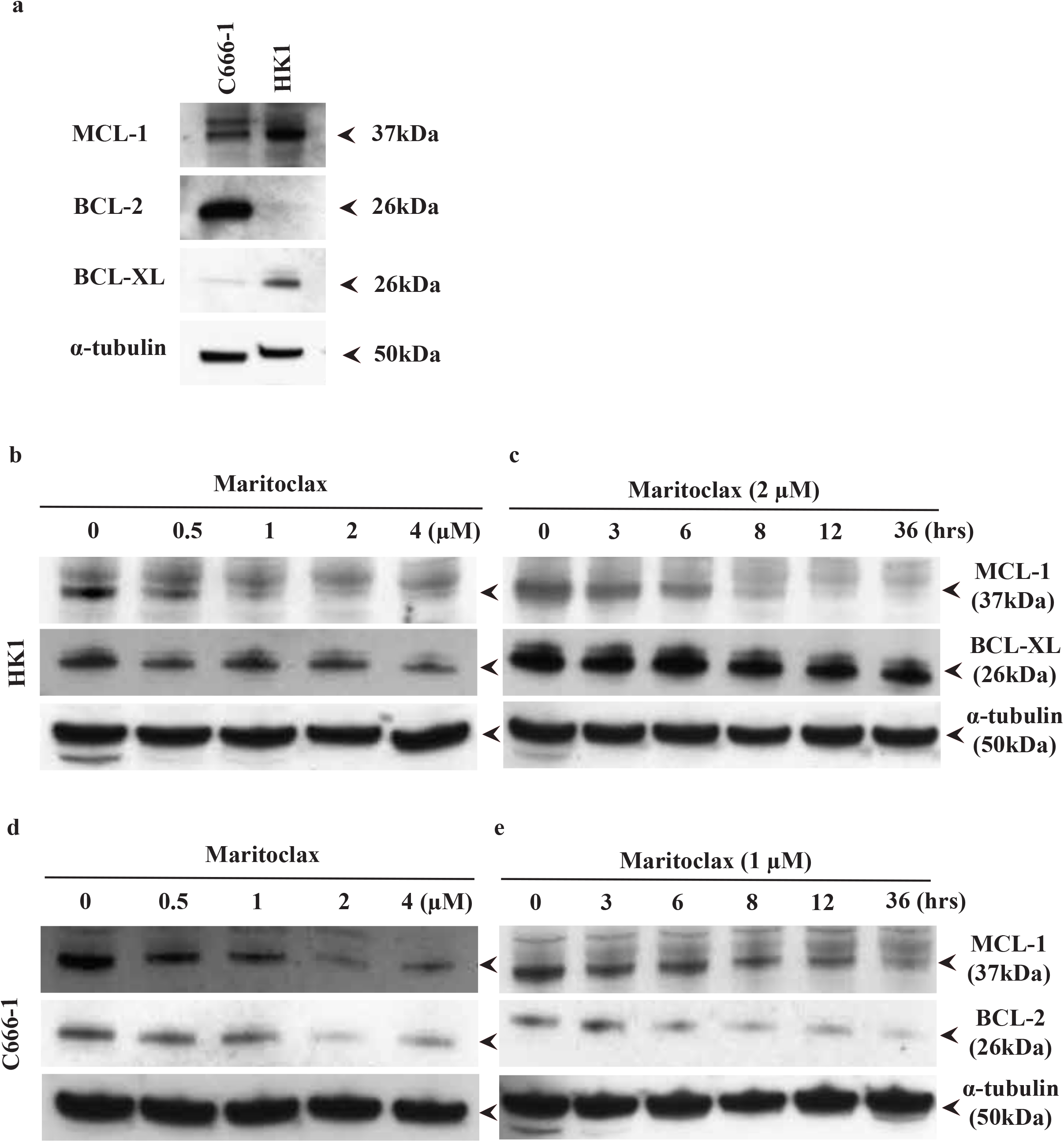
Maritoclax repressed the anti-apoptotic proteins in a dose- and time-dependent manner. (a); Basal expression levels of the anti-apoptotic proteins in the NPC cell lines. The basal expression levels of MCL-1, BCL-2 and BCL-XL in the C666-1 and the HK1 cells were determined by SDS-PAGE gel electrophoresis. NPC cell lines HK1 and C666-1 cells were treated with (b, d); escalating doses of Maritoclax for 24 hours to study the dose response or (c, e); with 2 μM or 1 μM of Maritoclax in the HK1 and C666-1 cells, respectively, at the indicated time points to study the time response. Immunoblot analyses illustrated that repression of MCL-1 by Maritoclax in both cell lines were dose- and time-dependent. (b-c); Maritoclax demonstrated a modest dose and time effect on the expression level of BCL-XL in the HK1 cells. (d-e); Maritoclax had a transient dose effect but a strong time effect on the level of BCL-2 in the C666-1 cells. Alpha-tubulin was used as the loading control. Arrows indicate the protein bands probed on the x-ray films.

Next, we determined the effect of Maritoclax on MCL-1 protein level. The HK1 and C666-1 cells were treated with Maritoclax at different doses and time points. Maritoclax reduced MCL-1 protein level in both cell lines at concentration as low as 0.5 μM and the level of MCL-1 continued to reduce as the drug concentration increased (Fig. 1b, d). Maritoclax also resulted in a time-dependent, reduction of MCL-1. In both cell lines reduction of MCL-1 was obvious at 8 hours and continued to reduce with time (Fig, 1c, e). Hence, the net effect of Maritoclax on MCL-1 protein level was both time- and dose-dependent. The effect of Maritoclax on the levels of BCL-XL and BCL-2 were also investigated. In the HK1 cells, the level of BCL-XL and not BCL-2 was investigated, as the basal level of BCL-2 in this cell line was too low (Fig. 1a) and *vice versa* in the C666-1 cells. Treatment of HK1 cells with Maritoclax repressed BCL-XL modestly in a dose- and time-dependent manner (Fig. 1b, c). Interestingly, in the C666-1 cells, increasing concentrations of Maritoclax exhibited a transient dose effect on the level of BCL-2 (Fig. 1d). The reduction of BCL-2 was near complete at 2 μM of Maritoclax but at 4 μM the protein level was restored (Fig. 1d). However, Maritoclax exhibited a strong time-effect on the level of BCL-2 in the C666-1 cells (Fig. 1e). The ability of Maritoclax to repress the other anti-apoptotic proteins indicates that it is not a selective MCL-1 inhibitor.

### NPC cells sensitive to single agent treatment of Maritoclax in a dose-dependant manner

The sensitivity of the HK1 and C666-1 cells were first tested to single agent Maritoclax and ABT-263 at increasing concentrations. Unexpectedly treatment with Maritoclax resulted in a profound shift of the dose-response curve to the left indicating a single agent activity of Maritoclax in inhibiting cell proliferation of both cell lines (Fig. 2a, b – open diamond). Given the single agent activity of Maritoclax, combination with ABT-263 at 1:1 drug concentration only resulted in a minimal shift of the dose-response curve in both cell lines (Fig. 2a, b – open square).

**Figure 2.**
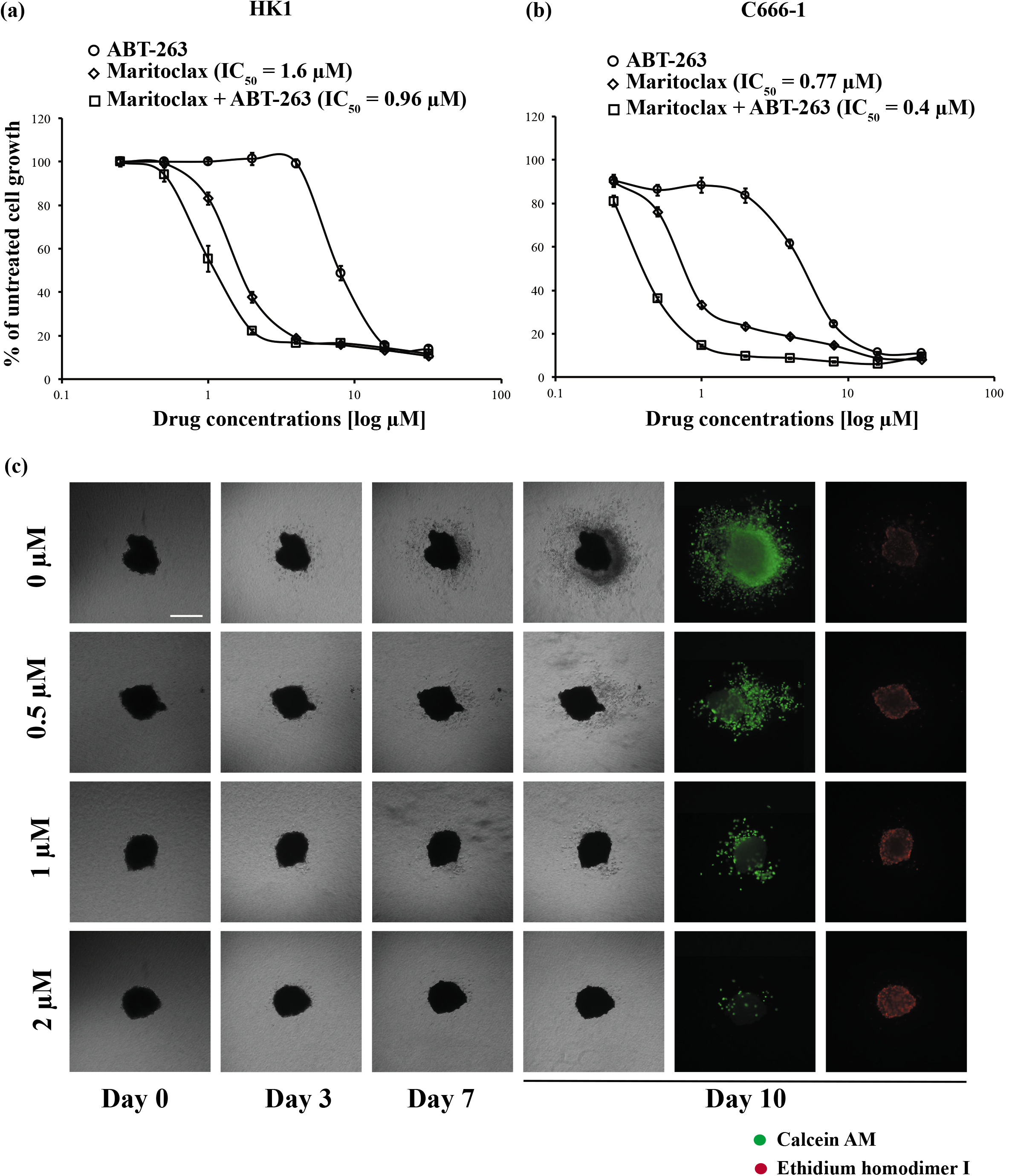
Maritoclax exhibited single agent activity in inhibiting cell proliferation of the NPC cells. (a); HK1 cells and (b); C666-1 cells were treated with increasing concentrations of ABT-263 (0-32 μM) (open circle) or Maritoclax (0-32 μM) (open diamond) or combination of ABT-263 and Maritoclax (open square) at 1:1 drug concentration ratio for 72 hours. Cell proliferation was assessed using the SyBr Green I assay. Points represent mean ± SEM of four experiments. (c); One representative experiment showing HK1 spheroids demonstrating decrease viable cells (calcein-AM) and increase dead cells (Ethidium homodimer I) after treatment with single agent Maritoclax at increasing concentrations over 10 days. Medium and drug were replenished every 72 hours. Size bar: 500 μM

Similar to the 2D findings, the spheroids established from the HK1 cells were sensitive to single-agent Maritoclax in 10-day assays, reflected in dose-dependent inhibition of invasion as well as decreased viable cell staining (Fig. 2c - calcein-AM staining) and increased dead cell staining (Fig. 2c - ethidium homodimer I). More notably there were no resistance cells proliferate and invade the matrix over 10 days.

### Maritoclax and ABT-263 synergistically inhibit cell proliferation

Next, we tested the sensitivity of the HK1 cells to combination of ABT-263 and Maritoclax. Considering the single agent activity of Maritoclax, we tested the sensitivity of cells to concentrations below the IC_50_ values of single agent Maritoclax combined with increasing concentrations of ABT-263. A fixed dose of Maritoclax (0.5 or 1 μM was added to escalating doses of ABT-263 (0-32 μM) (Fig. 3a). Concentration of Maritoclax at 0.5 μM exhibited only a 1.7-fold sensitization of the HK1 cells to ABT-263 but increased to 4-fold when concentration of Maritoclax was increased to 1 (Additional file 1). Strong synergism with multiple concentrations of ABT-263 was attainable at 1 μM Maritoclax (Fig. 3a). Given that the IC_50_ concentration of single agent Maritoclax on C666-1 cells was < 1 μM, combination with ABT-263 was not tested.

**Figure 3.**
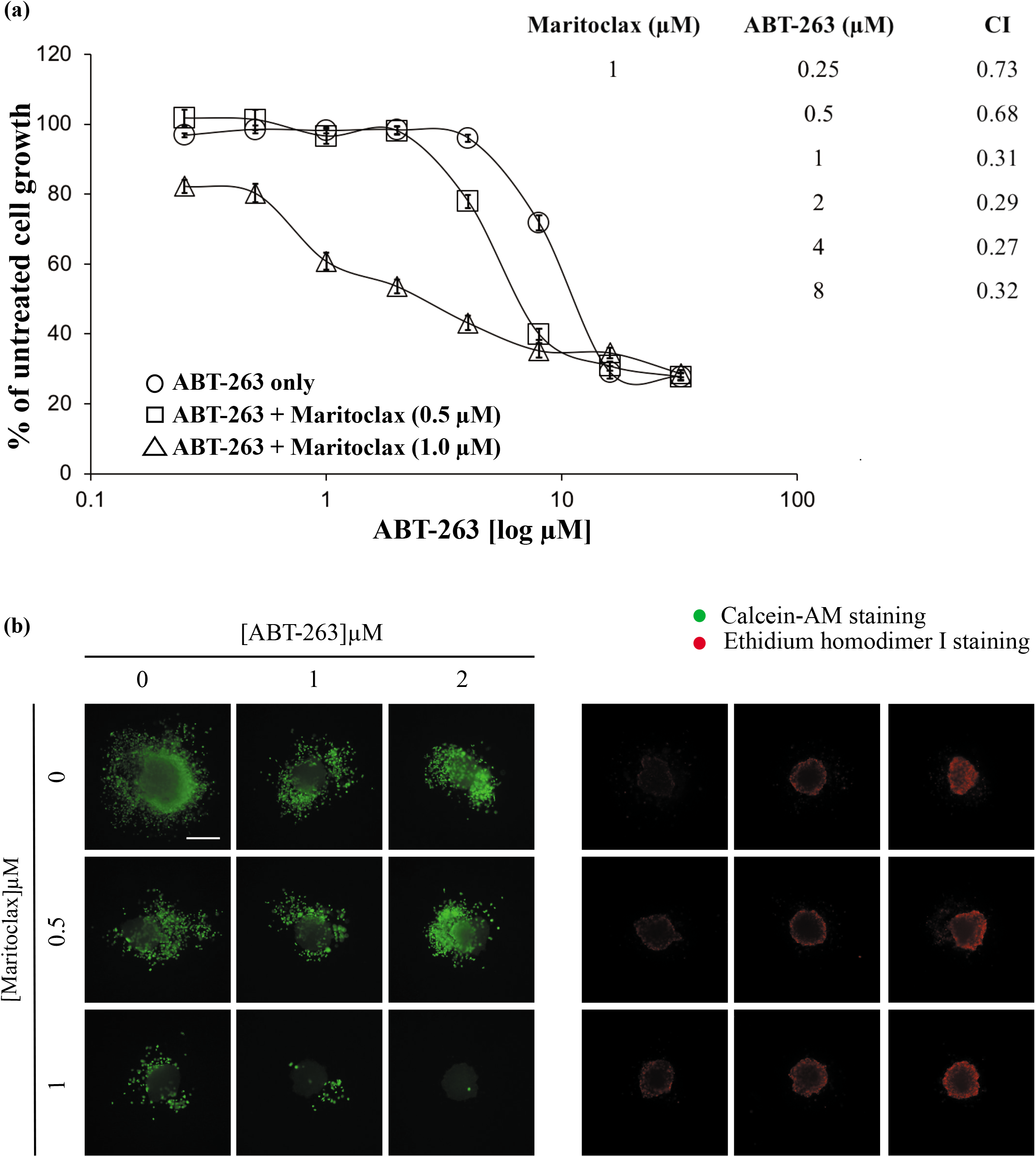
Synergistic anti-proliferative effects of ABT-263 and Maritoclax in the HK1 cells. (a); HK1 cells were treated with increasing concentrations of ABT-263 (0-32 μM) in the presence or absence of either 0.5 or 1 μM of Maritoclax for 72 hours. Cell proliferation was assessed using the SyBr Green I assay. Points represent mean ± SEM of four experiments. Combination index (CI) for the drug combination was calculated using the CalcuSyn 2.11 software (Biosoft Inc, Cambridge, UK). A CI of <1 indicates synergistic effect. (b); One representative experiment showing HK1 spheroids demonstrating decrease viable cells (calcein-AM) and increase dead cells (Ethidium homodimer I) after treatment with single agent Maritoclax and ABT-263 and combination of both drugs at increasing concentrations over 10 days. Size bar: 500 μM

The spheroids generated from the HK1 cells were also sensitized to ABT-263 by Maritoclax. The spheroids were treated with ABT-263 and Maritoclax, either alone or in combination over 10 days with medium and drugs replenishment every 72 hours. In the presence of 1 μM Maritoclax, there was obvious sensitization to ABT-263 reflected in dose-dependent inhibition of invasion. This also manifested as reduced cell viability (decreased Calcein AM staining and increased Ethidium homodimer I staining) (Fig. 3b).

## Discussion

Given that Maritoclax promoted proteasomal degradation of MCL-1 [9, 17], there was a reason to believe that combination with ABT-263 may overcome resistance and sensitize the NPC cells to ABT-263. Interestingly, a number of studies reported that Maritoclax did not display a stringent selective cytotoxicity on MCL-1 dependent cell lines [18, 19]. Human leukemia cell line RS4;11 which is dependent on BCL-2 for survival was more sensitive to Maritoclax treatment compared to HeLa cells which are dependent on MCL-1 for survival. Furthermore Maritoclax treatment did not alter the expression of MCL-1 in both these lines [18]. In another study, Maritoclax was shown to repress MCL-1 but this effect was not long lasting as the level of MCL-1 was restored at later time points in the MCL-1 dependent H460 cells. However, the drug did not repress the levels of anti-apoptotic proteins BCL-2, BCL-XL and BCL-w in the H460 cells. In contrast, Dinaciclib a broad spectrum CDK inhibitor which was reported to down-regulate MCL-1 exhibited a lasting time effect on the level of MCL-1 in the H460 cells [19]. The same study reported that the treatment with Maritoclax led to more death of the MCL-1 deficient MEFs compared to wild-type MEFs indicating that apoptosis can be triggered via other mechanisms [19]. In our hands, besides repressing MCL-1, Maritoclax exhibited a modest/transient dose effect on the levels of BCL-XL and BCL-2 in the NPC cell lines and modest time effect on the level of BCL-XL in the HK1 cells but a strong time-effect on the level of BCL-2 in the C666-1 cell line indicating that the effect of Maritoclax on the other anti-apoptotic proteins could be cell-type specific. The ability of Maritoclax to repress more than one anti-apoptotic protein may explain the single agent activity of Maritoclax in both the NPC cell lines as repression of MCL-1 and BCL-2/BCL-XL may be sufficient to reduce the threshold for apoptosis activation. Similar single agent activity of Maritoclax was reported in melanoma cells [20].

There were number of studies which reported synergy between Maritoclax and ABT-263 in leukemia and melanoma cells *in vitro* [9, 17, 20]. Treatment with suboptimal concentration of Maritoclax (2 μM) repressed MCL-1 and markedly sensitized melanoma cell line UACC903 to ABT-263 by ~23-fold. Sensitization of the melanoma cells was also evident in the 3D melanoma spheroids [20]. Combination of ABT-737 with sub-lethal dose of 2 or 2.5 μM Maritoclax in the K562 or Raji cells sensitized the cells to ABT-737 by ~60- and 2000-fold, respectively [9]. Similarly, two acute myeloid cells (AML) namely HL60 and KG1a, which were resistant to ABT-737 due to prolonged culture with the drug, were sensitized to ABT-737 after combination with sub-lethal dose of 2 or 1 μM Maritoclax, respectively [17]. In our study we had to be more cautious with the concentrations of Maritoclax used when combined with ABT-263, as Maritoclax was able to inhibit cell proliferation as single agent at very low concentrations (< 2 μM). Treatment with 1 μM of Maritoclax, sensitized the HK1 cells to ABT-263 by 4-fold. Similar findings were obtained in the 3D spheroids. However, given that Maritoclax exhibited single agent activity in both cell lines, it appears that treatment with Maritoclax alone will be sufficient to inhibit cell proliferation of NPC cells. Nevertheless, Maritoclax could be used as a sensitizer to ABT-263 or other drugs if appropriate drug concentrations are applied.

### Limitations

Given that Maritoclax inhibits BCL-2/BCL-XL, future work should investigate the mechanism behind the suppression of these two proteins by Maritoclax. The possibilities may be that Maritoclax impairs protein translation or transcription of these proteins. Future studies should also investigate the effect of single agent Maritoclax in preclinical models and testing of Maritoclax as a sensitizer to next generation BH3 mimetics and chemotherapeutic drugs for cancer management.

### List of Abbreviations

NPC: Nasopharyngeal carcinoma
2D: 2-dimensional
3D: 3-dimensional

## Declarations

### Ethics approval and consent to participate

Not applicable

### Availability of data and material

All data generated or analysed during this study are included in this published article [and its supplementary information files].

## Acknowledgements

We would like to thank the Director General of Health Malaysia for his permission to publish this article and the Director of the Institute for Medical Research for her support. We would like to thank Professor George S.W. Tsao from the School of Biomedical Sciences, The University of Hong Kong for providing the HK1 cell line and staff at the Molecular Pathology unit, Institute for Medical Research, KL, Malaysia for helping us with the STR profiling of the NPC cancer cell lines.

## Consent for publication

Not applicable

## Author’ contributions

BSXL and NM-K conceived and designed the study. BSXL carried out the study. BSXL and NM-K analyzed the data. NMK wrote the manuscript. K-WL and AS-BK revised the manuscript. All authors read and approved the final manuscript.

